# A passive upper-limb exoskeleton effectively reduces shoulder muscle activity over a large shoulder workspace

**DOI:** 10.1101/2025.02.26.640421

**Authors:** Leon Lauret, Brent James Raiteri, Paolo Tecchio, Daniel Hahn

**Affiliations:** Human Movement Science, Faculty of Sport Science, Ruhr University Bochum, Bochum, North Rhine-Westphalia, Germany; School of Human Movement and Nutrition Sciences, The University of Queensland, Brisbane, Queensland, Australia

**Keywords:** Musculoskeletal disorder, Overhead work, Exoskeleton, Joint angle, Subjective feedback

## Abstract

Industrial upper-limb exoskeletons are used to offload the upper limb during overhead tasks with the aim of preventing musculoskeletal disorders to the shoulder. Although numerous studies showed reduced shoulder muscle activity during upper-limb exoskeleton use for overhead postures, it remains unknown whether and how upper-limb exoskeletons provide support over a large shoulder workspace beyond overhead work. Therefore, this study evaluated the *Ottobock Paexo Shoulder* over a large shoulder workspace from overhead to hip height with shoulder abduction and adduction. Upper body kinematics, muscle activity, and subjective user feedback were obtained by three-dimensional motion capture, bipolar surface EMG and questionnaires, respectively, and captured while participants performed static and dynamic work tasks with an electric screwdriver. Participants completed these tasks while 1) not wearing the exoskeleton, 2) wearing a disengaged exoskeleton, with 3) moderate exoskeleton support, and with 4) high exoskeleton support. Exoskeleton support reduced deltoid muscle activity (-9 to -24s%, *p*≤.001) in postures with an abducted shoulder or extended elbow, including non-overhead postures. Exoskeleton support modestly decreased shoulder flexion (-3 to -5°, *p*≤.001) and increased shoulder abduction (2 to 5°, *p*≤.032), but the movement patterns during the dynamic task were unaffected. Additionally, exoskeleton-related effects increased with increasing support, but the subjective perception of change also increased, and perceived comfort decreased. Our results indicate that the tested exoskeleton provides support beyond overhead work and that there is a trade-off between exoskeleton support and subjective perception. Accordingly, further optimization of user-exoskeleton interaction is warranted for long-term prevention of musculoskeletal disorders in overhead workers.

## Introduction

Musculoskeletal disorders (MSDs) to the shoulder are a leading cause of work-related sick leave in overhead workers (Barthelme et al., 2021) and substantially affect worker well-being and productivity (German Federal Ministry of Labor and Social Affairs, 2019). Thus, the industrial sector has a great interest in reducing the risk of shoulder MSDs in overhead workers. One potential solution to reduce shoulder MSD risk is to offload one or both shoulders with a passive upper-limb exoskeleton.

Research evaluating upper-limb exoskeletons found reduced muscle activity of various shoulder and back muscles while performing overhead drilling and wiring tasks with shoulder flexion angles exceeding 90° (De Vries et al., 2019; Hefferle, Snell & Kluth, 2021; Kim et al., 2018; Otten, Weidner & Argubi-Wollesen, 2018; Schmalz et al., 2019; Van Engelhoven et al., 2018). These muscle activity reductions occurred alongside reduced ratings of exertion during exoskeleton use (De Vries et al., 2019; Maurice et al., 2019). However, such findings are limited to overhead work.

Accordingly, it remains unknown whether typical upper-limb exoskeletons provide support over a large shoulder workspace including in non-overhead working postures (De Bock et al., 2021; De Vries & De Looze, 2019; Kim et al., 2021). Contrary to a potential support, upper-limb exoskeletons might oppose non-overhead work. This is because typical passive exoskeleton mechanisms are based on springs or elastomers; when the exoskeleton user raises their arms, the exoskeleton’s springs or elastomers release stored energy to return to their resting length, which reduces the shoulder torque that needs to be generated by active muscle contraction to maintain an overhead posture. However, when exoskeleton users lower their arms, the same springs or elastomers are stretched and thus provide resistance to the intended movement. Consequently, the typical passive exoskeleton mechanism might limit the beneficial effects of upper-limb exoskeletons over a large shoulder workspace (De Vries & De Looze, 2019; Van Engelhoven & Kazerooni, 2019).

The exoskeleton itself might also increase the activity of back muscles during overhead work. This is because upper-limb exoskeletons typically transfer load away from the shoulders and towards the core via a frame connected to the upper arms, which offloads the shoulders. However, the load transfer to the core could result in adverse effects by increasing activity in other muscles such as the erector spinae or latissimus dorsi (De Bock et al., 2021, Schmalz et al., 2019). Further, non-overhead postures and downward movements might require additional back muscle activity for stretching the exoskeleton’s springs or elastomers (De Vries & De Looze, 2019).

The exoskeleton’s effects in different working postures are likely to be mediated by the level of exoskeleton support, but the interaction between posture and exoskeleton support has received little attention to date, despite the importance of this knowledge for optimizing the beneficial effects of exoskeleton use. Most studies on upper-limb exoskeletons simply compared effects on tasks performed with and without an exoskeleton. Further, inconsistent definitions of ‘optimal’ support levels have been used; for example, Maurice et al. (2019) fully compensated the arm’s weight with the shoulder and elbow at 90° flexion, whereas Schmalz et al. (2019) compensated for only 70% of arm weight at 105° arm elevation. Consequently, there is a need to test different levels of support during upper-limb exoskeleton use and to evaluate the effects in different working postures.

Therefore, the aims of this study were: A) to quantify the effects of an industrial upper-limb exoskeleton over a large shoulder workspace, and; B) to evaluate how different exoskeleton support levels affect a dynamic work task over a large shoulder workspace. We hypothesized that A1) in line with previous research, exoskeleton support would offload the shoulder and reduce shoulder muscle activities in abducted and flexed arm postures only, and A2) that exoskeleton support would affect shoulder kinematics because of the exoskeleton’s shoulder support mechanism. We also hypothesized that B1) increased exoskeleton support would reduce shoulder muscle activities progressively more in arm postures at shoulder height and above, but increase back muscle activities in arm postures below shoulder height. Lastly, we hypothesized that B2) the subjective feedback to different exoskeleton support would not necessarily become more positive as the objective exoskeleton-related benefits increased, making user feedback essential when deciding on the optimal level of support.

## Methods

### Participants

Eighteen healthy participants (7 women) participated in the study after providing written informed consent. The participants were physically active and had no surgery and no major or minor injury to the upper limbs prior to testing (<24 months). Participants (16 right-handed, 2 left-handed) had a mean (standard deviation) age of 26.8 (1.7) years (range: 24.8-32.0 years), height of 181.0 (11.0) cm (range: 162.0-195.0 cm), and mass of 76.1 (13.3) kg (range: 58.0-94.6 kg). All participants were university students with no industrial experience or familiarity with exoskeleton use. The experimental protocol and procedures were approved by the Ethics Committee of the Faculty of Sport Science at Ruhr University Bochum (EKS V 17/2022) and conducted in accordance with the Declaration of Helsinki.

### Experimental setup

This study evaluated the effects of the *Ottobock Paexo Shoulder* exoskeleton using two standardized screw driving tasks with an electric screwdriver. To standardize the tasks, a 1×1 meter board of adjustable height was fixed to a rack (Fig. 1A). A matrix of screws was installed on the board with a screw-to-screw distance of 16 cm (6×6 matrix, Fig. 1B) or 40 cm (3×3 matrix, Fig. 1C), respectively. The two different matrices were used to optimize the assessment of the entire shoulder workspace; the 3×3 matrix was used for the static task to test endpoints of shoulder flexion/extension and abduction/adduction, whereas the 6×6 matrix was used for the dynamic task to provide a more comprehensive coverage of the workspace and movement patterns during continuous motion across screw locations. The screws (M6, 70×5mm) had to be driven in using a 1.5kg electric screwdriver (Festool GmbH, Wendlingen a.N., Germany), which took 10 s per screw on average. The height of the task board was set by aligning the acromion process of the participants’ dominant arm to a predefined position between screws B3 and B4 (red cross; Fig. 1). Afterwards, the participants stood at a comfortable position away from the board; the distance was then marked on the floor using tape and the participants returned to this position for each subsequent trial.

**Figure 1.**
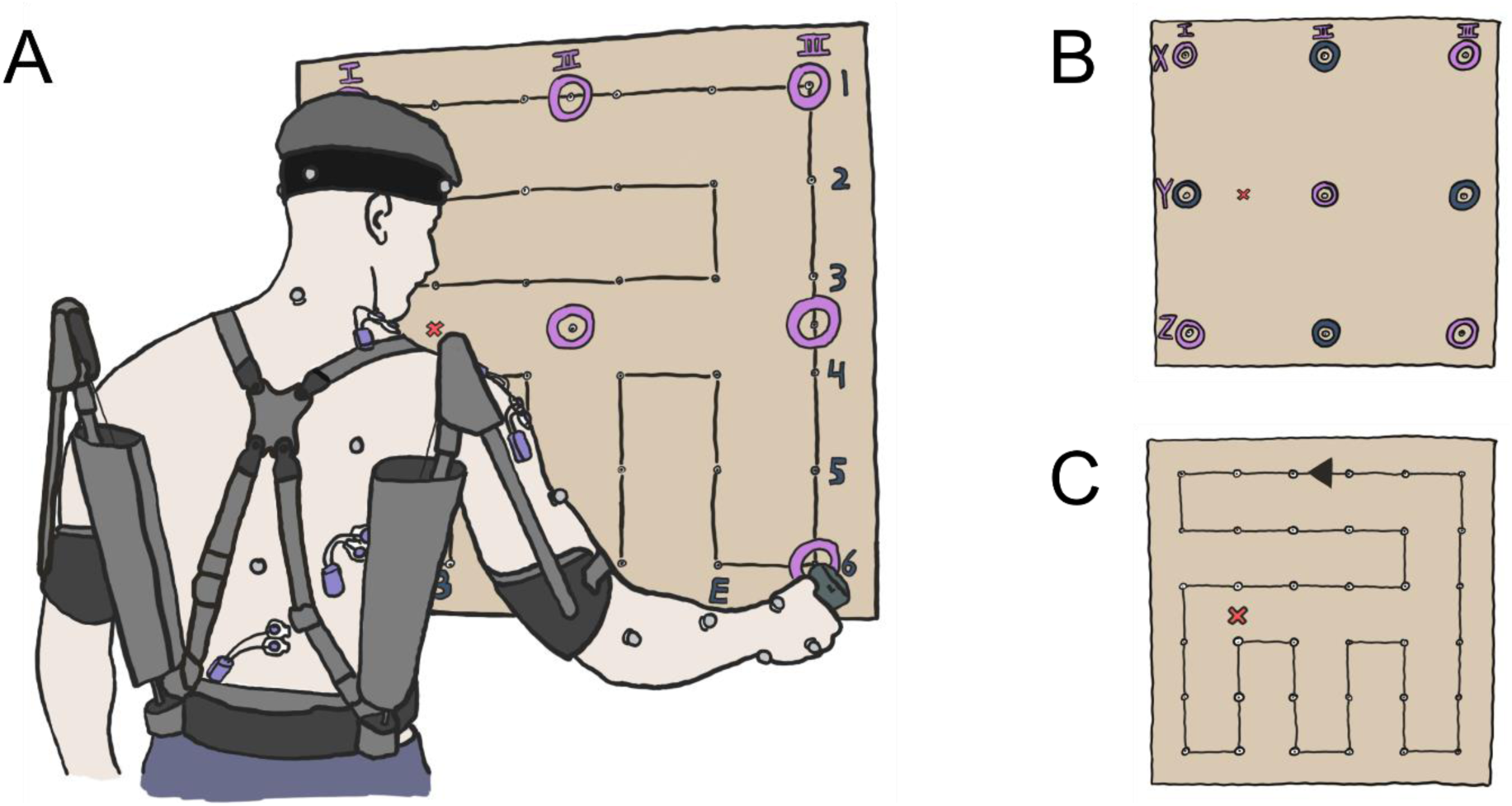
Illustration of the experimental setup. A) A wooden board with a 6×6 matrix of screws was used for two tasks, which were performed for each of the four support levels. The participants’ acromion process (dominant arm) was aligned to a reference point (red cross) and upper body kinematics were recorded through the use of sixteen reflective markers, placed according to a modified version of Vicon’s upper-body plug-in gait model. Surface electrodes were placed on upper trapezius (TRAP), anterior deltoid (AD), medial deltoid (MD), latissimus dorsi (LAT) and erector spinae (ES). B) In a static work task (STA), joint kinematics and muscle activities were recorded during the individual driving phase of nine screws (grey dots), and recordings from five screw locations (purple circles) were used in the statistical analysis. C) dynamic work task (DYN) consisted of drilling in the entire 6×6 matrix of screws in a predefined pattern (black line), which took around six minutes to complete. The arrow indicates the movement direction between screws, and the predefined pattern finished at the same randomized screw that the participants started at.

### Exoskeleton support setup

The manufacturer’s recommendation for an optimal support level (i.e. *“The level of support is set correctly when the arms drop down due to gravity alone, with no additional force. The level of support is not permitted to exceed this value, but can be freely chosen below this”;* Ottobock SE & Co. KGaA, 2019, p.24) lacks specificity and does not give clear guidance for practical implementation. To address this uncertainty in setting the level of support, we defined and tested four support levels and systematically evaluated their effects on upper body kinematics and muscle activity levels. The four support levels included: 1) no exoskeleton use (NoEXO), which acted as a reference condition to help evaluate the effects of exoskeleton use during each working task (see next paragraph) and 2-4) exoskeleton use during the same two working tasks, with three support levels: 2) no support with the elastomers disengaged (EXOdis), 3) moderate support (EXOmod), and 4) high support (EXOhigh). EXOdis served to evaluate effects from just wearing the exoskeleton during the working tasks. EXOhigh was designed to support the participant’s estimated arm weight at 90° of shoulder abduction, with arm weight estimation based on De Leva (1996) and defined as 4.49% or 4.94% of the participant’s body mass for women and men, respectively. However, because the maximal support of the exoskeleton was limited, the estimated arm weight of the men exceeded the maximal exoskeleton support. Consequently, the differences between support levels varied between men, and from our sample of women and men, the mean EXOhigh support was only 90% of the estimated arm weight (range: 75%-100%). We decided not to choose a lower, standardized support level for the EXOhigh condition because of the difficulty in determining the maximal available support without a-priori knowledge of our participants’ characteristics, and because we also wanted to test the limits of exoskeleton support to investigate whether this would result in adverse effects. The support provided by the exoskeleton during EXOmod was simply half of that provided during EXOhigh. The mean EXOmod support for our sample was 45% (range: 38-50%) of the estimated arm weight. The order of exoskeleton conditions was randomized for each participant.

### Working tasks

For each support level, two standardized working tasks were performed with a rest period of five minutes between tasks. The first task was designed to evaluate the exoskeleton’s effects in nine different arm postures. A 3×3 matrix of screws were individually driven in for ∼10 s each with 30 s of rest (i.e. standing with arms by sides) in between every screw (static task, STA; Fig. 1B). Kinematics and muscle activities were recorded for each screw, and the order of screws was randomly assigned for each condition and participant.

The second task was designed to evaluate the exoskeleton’s effect during a dynamic task (DYN) over a large shoulder workspace and consisted of driving in 36 screws of a 6×6 matrix for ∼10 s each without a break in between screws. After each screw, the participants moved onto the next screw in a predefined loop pattern (Fig. 1C). The starting screw was randomly assigned to each participant in the first condition and set among conditions. The randomization was performed in this way to limit fatigue and learning effects among support levels without changing the demands of the loop pattern in terms of horizontal and vertical distance covered. To quantify the fatigue accumulated during DYN, the loop-pattern finished with driving in the starting screw a second time, which resulted in 37 screws being driven in by the participants over six minutes. During the DYN task, kinematics and muscle activities were recorded for the first five and last five screws among the four support levels. After DYN, participants remained seated for fifteen minutes and during this period filled out a questionnaire if the exoskeleton was used (see *Subjective feedback*).

### Rest between support levels

Before each STA and DYN task commenced with a new support level, a reference screw (centre screw Y-II, Fig. 1B) was driven in without the exoskeleton for 10 seconds to assess fatigue throughout the experiment. If the muscle activity level of the anterior deltoid exceeded the muscle activity level during the very first reference screw recording by more than 5%, three minutes of additional rest was provided and this process was repeated until this criterion was met. Muscle activity was determined as a centered moving root-mean-square (RMS) amplitude with a window duration of 2 s.

### Muscle activity

Muscle activity levels were obtained from the anterior deltoid (AD), medial deltoid (MD) and upper trapezius (TRAP) (Fig. 1A) via bipolar surface electromyography (EMG) using a wireless Myon system (myon AG, Schwarzenberg, Switzerland) and 24 mm bipolar Ag/AgCl electrodes (Cardinal Health 200, LLC, Waukegan, USA). During DYN, muscle activity levels of the latissimus dorsi (LAT) and erector spinae (ES) were also recorded. Electrodes were only placed over muscles on the dominant hand’s side of the body as participants performed both working tasks with their dominant hand. Surface EMG signals were recorded at 1000 Hz using Vicon Nexus (Vicon Nexus 2.12, Vicon Motion Systems Ltd. Oxford, UK) and electrodes were placed with an inter-electrode distance of 2 cm following SENIAM recommendations (Hermens et al., 1999). To improve signal quality, the skin was first shaved (Teqler, Wecker, Luxembourg), exfoliated (Nuprep, Weaver and Company, Aurora, United States), and then disinfected (alcohol-based hand disinfectant; Sterillium, Hartmann, Heidenheim, Germany). The electrodes and corresponding wireless sensors were taped to the skin using kinesio tape (MIKROS GmbH, Hamburg, Germany).

### Kinematics

Upper limb and upper body kinematics were recorded using a Vicon motion capture system. Eight infrared cameras (Vantage V5, 5 Megapixels) recorded sixteen (for right-handed participants; for left-handed participants, the right back marker was excluded) reflective markers (14 mm diameter) at 100 Hz. The markers were located on specific anatomical landmarks on the head, shoulder, elbow, wrist and back (Fig. 1A) according to a modified marker set based on the upper-body plug-in gait model (Vicon Motion Systems, Plug-in Gait Reference Guide, 2023) without markers on the non-dominant shoulder and arm.

### Subjective feedback

A questionnaire was used to evaluate the participants’ perception of physical demand, perceived change, and comfort during exoskeleton use (for further details, see supplementary material). The participants’ answers were recorded on a scale derived from the Comfort rating scales (CRS; Knight et al., 2002) using 21 points, which were scored from 0 (“low”) on the left to 20 (“high”) on the right. The questionnaire was completed after task completion of STA and DYN with one exoskeleton support level. This allowed us to assess differences in subjective feedback for the different exoskeleton support levels (i.e. EXOdis, EXOmod, and EXOhigh).

### Data processing and analysis

Data analysis was performed using custom-written MATLAB scripts (R2021a, 64-bit version; The MathWorks Inc., Natick, Massachusetts, USA). EMG signals were bandpass-filtered at 20–250 Hz (based on visual checks of fast Fourier transforms) with a zero-lag fourth-order Butterworth filter. To maximize the number of EMG channels without motion artifacts, each recording was then split into overlapping bins of 1 second and the bin with the minimum energy content was identified. The mean EMG amplitude of each muscle’s minimum energy content bin was then calculated using a DC offset remove followed by a 1-s window RMS amplitude. The differences in RMS amplitude among exoskeleton support levels relative to NoEXO were then calculated for STA using the symmetrized percent difference (s%, Nuzzo, 2018). The s% values were used to construct muscle activity level heatmaps for the 3×3 matrix of screws, and these heatmaps were combined with information on the shoulder and elbow joint angles for each screw (Fig. 2&3). For DYN, the same calculations were made at the start (reference) and end of the task for the same screw.

**Figure 2.**
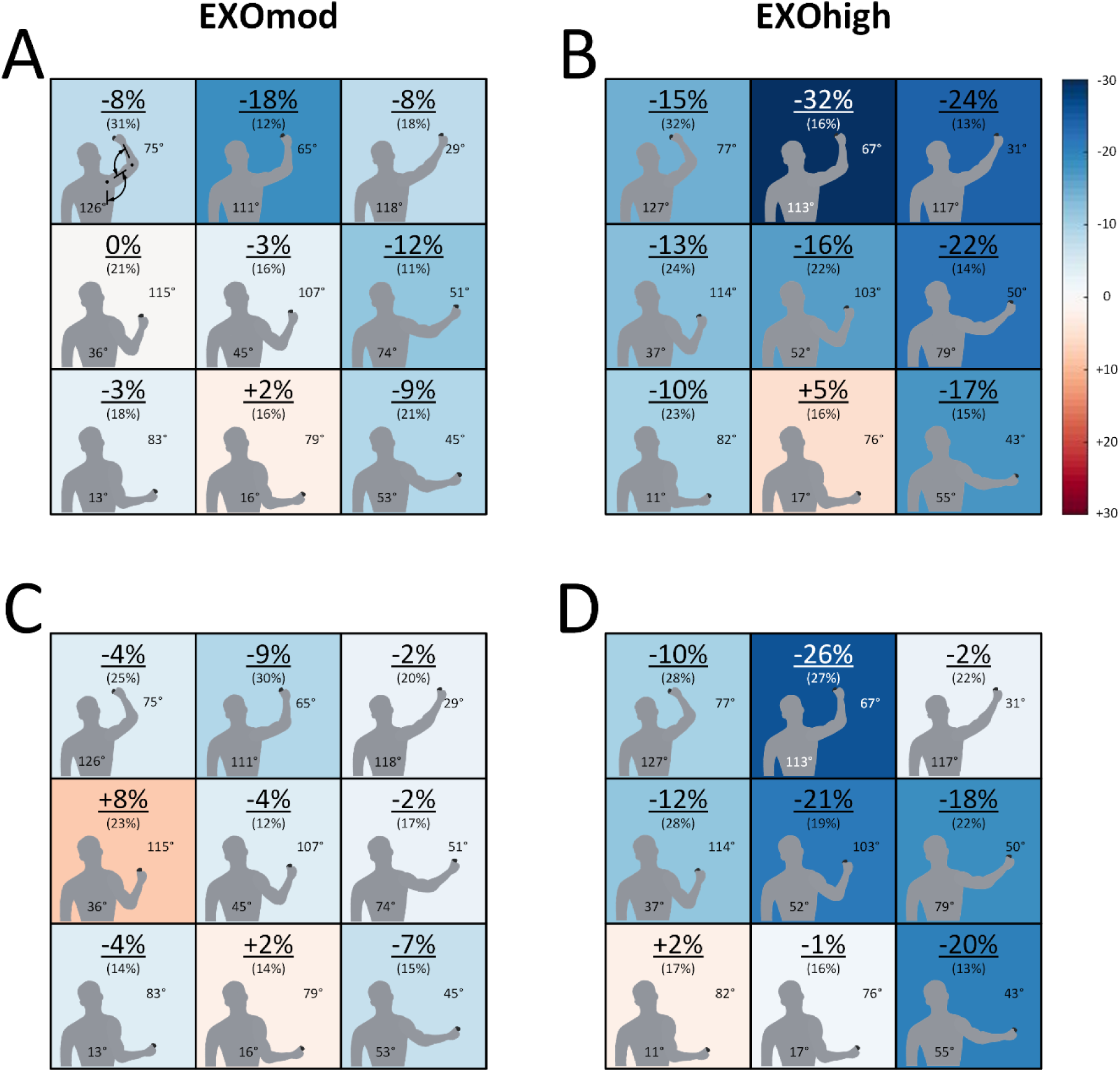
**Heatmaps (n = 17) in the EXOmod (A&C) and EXOhigh (B&D)** conditions showing the mean (SD) differences in anterior deltoid (A&B) and medial deltoid (C&D) muscle activity level relative to NoEXO at different screw locations during STA. Muscle activity level differences were calculated as symmetrized percent differences, and shoulder abduction and elbow flexion angles are shown at each screw location. The heatmaps show that exoskeleton support reduced deltoid muscle activity levels in arm postures away from the core, but the effects were limited in arm postures close to the core. Higher support (B&D vs. A&C) led to a larger overall reduction in deltoid muscle activity levels over the entire workspace.

Shoulder and elbow joint angles were calculated from the 3D motion capture recordings using built-in pipelines in Vicon Nexus. In cases of missing or obstructed markers, manual reconstruction was performed using open-source MATLAB scripts (www.vicon.com, see supplementary material). Afterwards, the mean shoulder flexion (SHOflex), abduction (SHOabd) and internal rotation (SHOrot), as well as the elbow flexion (ELBflex) angles were determined for each individual screw during STA. For DYN, joint angles were determined for the first (START) and identical last screw (END). START and END were identified by subjective identification of the start and end of the constant velocity phase of the finger marker, corresponding to the time spent driving a screw (for further details, see supplementary material). To evaluate how the participants movements were affected by exoskeleton use during DYN, movement pathlength of the finger marker was calculated for the first five and last five screws. The pathlength was calculated from the x, y, and z marker coordinates using the following equation:

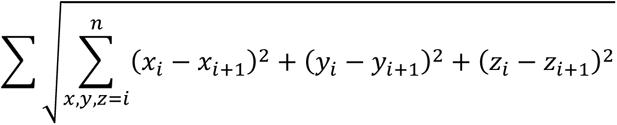

### Statistical analysis

To reduce the number of comparisons done for STA, only data from the outer screws and neutral middle screw were considered during statistical analysis (Fig. 1B). Statistical analysis was performed using GraphPad Prism (9.1.2 64-bit version, San Diego, California, USA) and α = 0.05 (two-tailed). Two-way repeated-measures mixed-effects models with the Greenhouse-Geisser correction were used to determine the effects of support level and working posture (i.e. screw location) on muscle activity level (AD, MD or TRAP) and joint angle (SHOflex, SHOabd, SHOrot or ELBflex) during STA. The same model was also used to determine the effects of support level and time on muscle activity levels (AD, MD, TRAP, LAT or ES), joint angles (SHOflex, SHOabd, SHOrot or ELBflex), and pathlength during DYN. Following a significant main effect or interaction, Holm-Šídák post-hoc comparisons were performed (Holm, 1979) to identify which exoskeleton conditions were significantly different to NoEXO. Friedman tests were performed on the questionnaire data and following a significant main effect, Dunn’s tests were performed (Dunn, 1964). Data are reported as mean (standard deviation) unless indicated otherwise.

## Results

### Exclusions

One participant’s dataset was excluded because of motion artifacts in the EMG signals, which resulted in *n* = 17.

### Static task

#### Muscle activity

Muscle activities of AD and MD were affected by support level (*p*<.001) and posture (*p*<.001), but there was no significant interaction between support level and posture (*p*=.069 and *p*=.083, respectively) (Fig. 2). Muscle activity of TRAP was also affected by support level (*p*=.001) and posture (*p*<.001), and there was a statistically significant interaction between support level and posture (*p*=.005) (Fig. 3).

**Figure 3.**
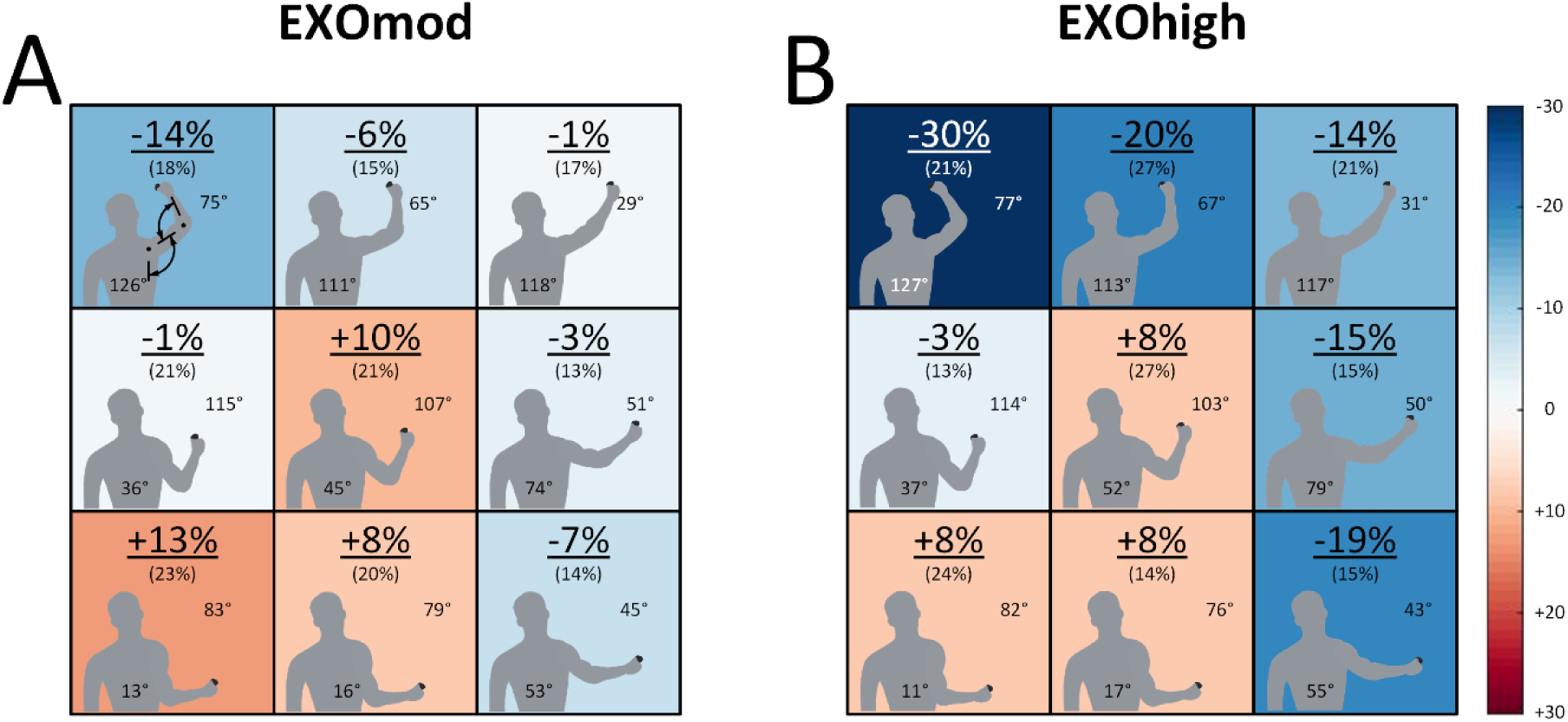
**Heatmaps (n = 17) in the EXOmod (A) and EXOhigh (B) conditions** showing the mean (SD) differences in upper trapezius muscle activity level relative to NoEXO at different screw locations during STA. Muscle activity level differences were calculated as symmetrized percent differences and shoulder abduction and elbow flexion angles are shown at each screw location. The heatmaps show that exoskeleton support reduced trapezius muscle activity levels in arm postures away from the core, but the effects were limited in arm postures close to the core. Higher support (B&D vs. A&C) led to a larger overall reduction in trapezius muscle activity levels over the entire workspace.

#### Kinematics

All shoulder angles (i.e. SHOflex, SHOabd, and SHOrot) showed significant interaction effects between support level and posture (*p*<.001). Compared with NoEXO, mean SHOflex was reduced in EXOdis (−3.61°, *p*<.001), EXOmod (−2.8°, *p*=.001), and EXOhigh (−5.0°, *p*<.001; Tab. 1). On the contrary, mean SHOabd increased with exoskeleton support in EXOdis (2.1°, *p*=.032), EXOmod (3.9°, *p*<.001), and EXOhigh (4.7°, *p*<.001; Tab. 2). Similarly, SHOrot showed a main effect of support level (*p*<.001) in EXOdis (−4.4°, *p* = .002), EXOmod (−5.8°, *p*=.015), and EXOhigh (−11.9°, *p* = <.001), as well as a main effect of posture (*p*<.001). ELBflex showed a significant interaction effect between support level and posture (*p*<.001), and there was a main effect of posture (*p* = <.001), but not support (*p* = .699).

**Table 1.** Mean (SD) shoulder flexion angles (n = 17) and mean differences between exoskeleton support levels compared with NoEXO during STA. Statistical analysis was performed on the corner screws and the middle screw (highlighted in black). Exoskeleton support reduced shoulder flexion angles in arm postures away from the core only. *indicates significant differences relative to NoEXO.

| Support level | Mean (SD) | Mean diff. | Mean (SD) | Mean diff. | Mean (SD) | Mean diff. |
| --- | --- | --- | --- | --- | --- | --- |
|  | XI |  | XII |  | XIII |  |
| NoEXO | 42 (14) |  | 30 (10) |  | 27 (6) |  |
| EXOdis | 35 (14) | -7 * | 22 (10) | -8 | 22 (7) | -5 * |
| EXOmod | 34 (17) | -7 * | 22 (11) | -8 | 22 (7) | -5 * |
| EXOhigh | 34 (16) | -8 * | 23 (10) | -7 | 15 (10) | -12 * |
|  | YI |  | YII |  | YIII |  |
| NoEXO | 27 (11) |  | 12 (9) |  | 22 (7) |  |
| EXOdis | 26 (10) | -1 | 10 (8) | -2 | 16 (6) | -6 |
| EXOmod | 31 (16) | 4 | 11 (9) | -1 | 17 (6) | -5 |
| EXOhigh | 30 (14) | 3 | 11 (9) | -1 | 19 (9) | -3 |
|  | ZI |  | ZII |  | ZIII |  |
| NoEXO | -4 (8) |  | -7 (8) |  | 17 (9) |  |
| EXOdis | -4 (8) | 0 | -6 (7) | 1 | 12 (9) | -5 * |
| EXOmod | -3 (6) | 1 | -6 (7) | 1 | 13 (10) | -4 * |
| EXOhigh | -5 (6) | -1 | -6 (6) | 1 | 13 (8) | -4 * |

**Notes.**
\* Within the screw, joint angle means compared to NoEXO differ ( $p < 0.05$ )

**Table 2.**
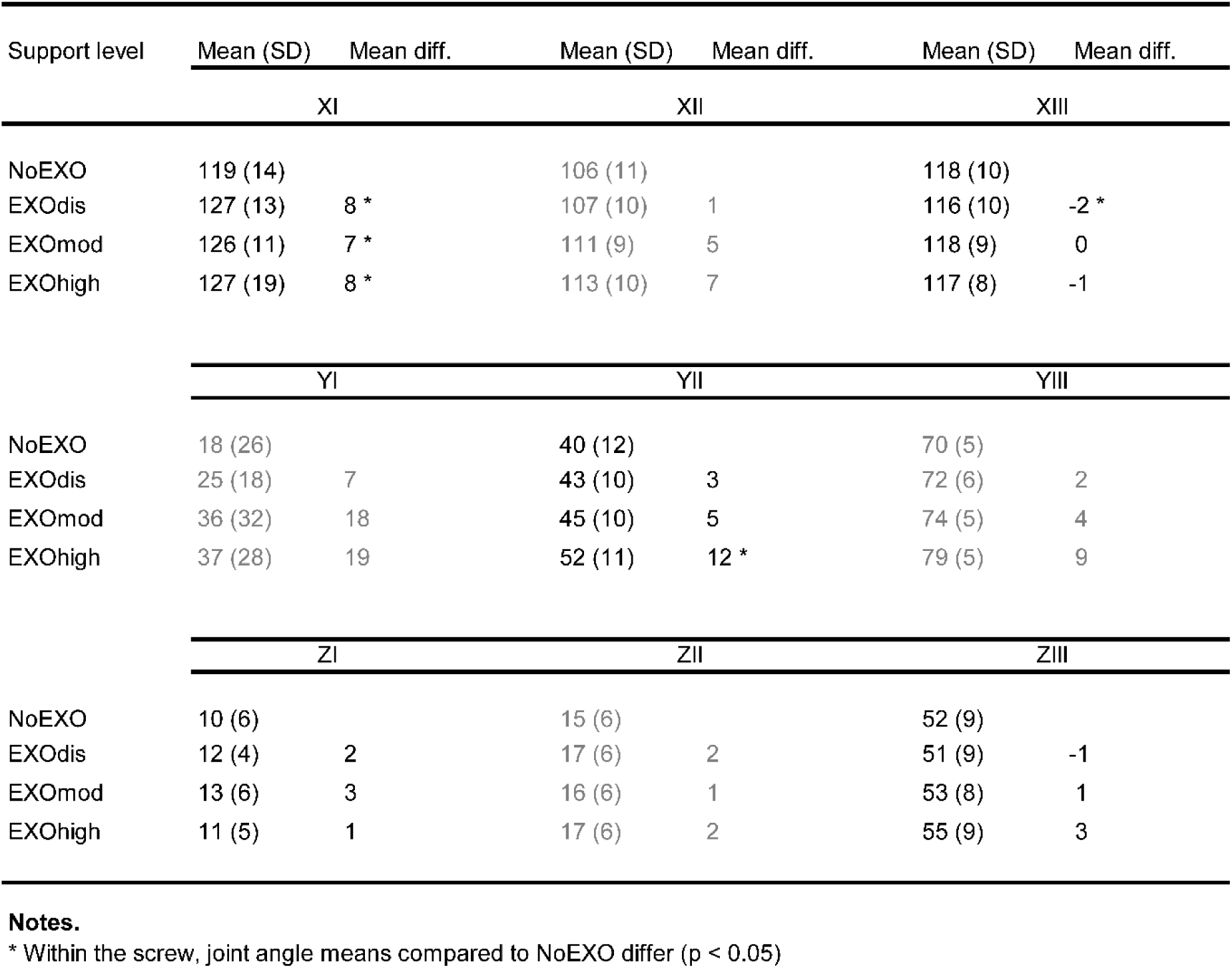
Mean (SD) shoulder abduction angles (SD) (n = 17) and mean differences between exoskeleton support levels compared with NoEXO during STA. Statistical analysis was performed on the corner screws and the middle screw (highlighted in black). Exoskeleton support increased shoulder abduction angles in positions with presumed lower shoulder torques, as increased shoulder abduction with increasing exoskeleton support is evident in positions where the shoulder was abducted, but the elbow remained flexed. *indicates significant differences relative to NoEXO.

| Support level | Mean (SD) | Mean diff. | Mean (SD) | Mean diff. | Mean (SD) | Mean diff. |
| --- | --- | --- | --- | --- | --- | --- |
|  | XI |  | XII |  | XIII |  |
| NoEXO | 119 (14) |  | 106 (11) |  | 118 (10) |  |
| EXOdis | 127 (13) | 8 * | 107 (10) | 1 | 116 (10) | -2 * |
| EXOmod | 126 (11) | 7 * | 111 (9) | 5 | 118 (9) | 0 |
| EXOhigh | 127 (19) | 8 * | 113 (10) | 7 | 117 (8) | -1 |
|  | YI |  | YII |  | YIII |  |
| NoEXO | 18 (26) |  | 40 (12) |  | 70 (5) |  |
| EXOdis | 25 (18) | 7 | 43 (10) | 3 | 72 (6) | 2 |
| EXOmod | 36 (32) | 18 | 45 (10) | 5 | 74 (5) | 4 |
| EXOhigh | 37 (28) | 19 | 52 (11) | 12 * | 79 (5) | 9 |
|  | ZI |  | ZII |  | ZIII |  |
| NoEXO | 10 (6) |  | 15 (6) |  | 52 (9) |  |
| EXOdis | 12 (4) | 2 | 17 (6) | 2 | 51 (9) | -1 |
| EXOmod | 13 (6) | 3 | 16 (6) | 1 | 53 (8) | 1 |
| EXOhigh | 11 (5) | 1 | 17 (6) | 2 | 55 (9) | 3 |

**Notes.**
\* Within the screw, joint angle means compared to NoEXO differ ( $p < 0.05$ )

### Dynamic task

#### Muscle activity

Muscle activities of AD and MD showed significant differences among support levels when compared between START and END (*p*<.001 for AD and MD), but there was no main effect of time (AD: *p*=.217; MD: *p*=.311), and no interaction between support level and time (AD: *p*=.557; MD: *p*=.164). Relative to NoEXO, muscle activities were lower in EXOhigh for AD (−25.4 s%, *p*<.001) and MD (−23.7 s%, *p*<.001), but were no significant relative differences in EXOmod or EXOdis (Fig. 4). TRAP muscle activity was not significantly affected by support level (*p*=.141) or time (*p*=.604), and there was no interaction between support level and time (*p*=.456). Similar results were found for LAT and ES muscle activities (LAT: support [*p*=.057], time [*p*=.549], interaction [*p*=.409]; ES: support [*p*=.242], time [*p*=.285], interaction [*p*=.255]).

**Figure 4.**
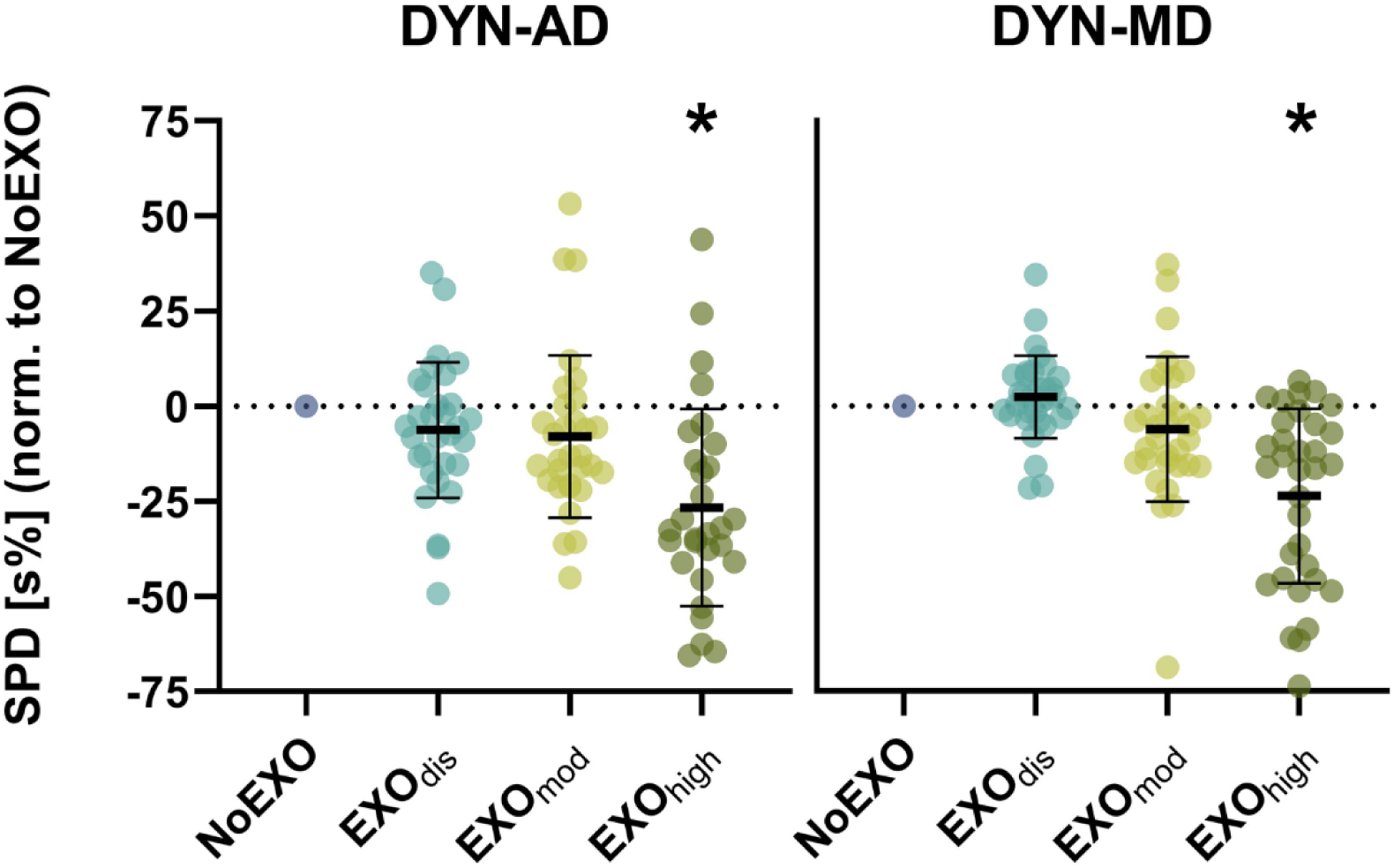
Symmetrized percent differences (SPDs) in anterior deltoid (AD) and medial deltoid (MD) muscle activities during DYN. Statistical analysis revealed significant differences among support levels without an effect of time. *indicates significant differences relative to NoEXO (mean differences = AD: −25.4 s%, p<.001; MD: −23.7 s%, p<.001).

#### Kinematics

Support level significantly affected SHOabd (*p*<.001) and SHOrot (*p*=.008) during DYN, but joint flexion (i.e. SHOflex and ELBflex) was unaffected by support level during DYN. Pathlength was only significantly affected by time (*p*=.013, support [*p*=.500], interaction [*p*=.388]), with pathlength increasing by 10% from START to END (2662 (93) mm vs. 2923 (81) mm). All mean joint angles for shoulder flexion, abduction, rotation and elbow flexion across at START and END during DYN are presented in Table 3.

**Table. 3.** Mean (SD) joint angles (°) for shoulder flexion, abduction, rotation and elbow flexion across at START and END during DYN. *indicates significant differences at the respective time point (i.e. START or END) relative to NoEXO.

| Support level | Shoulder flexion |  | Shoulder abduction |  | Shoulder rotation |  | Elbow flexion |  |
| --- | --- | --- | --- | --- | --- | --- | --- | --- |
|  | START | END | START | END | START | END | START | END |
| NoEXO | 15 (20) | 18 (19) | 58 (40) | 59 (45) | -29 (39) | -25 (49) | 89 (25) | 87 (30) |
| EXOdis | 12 (16) | 14 (17) | 60 (38) | 60 (43) | -25 (39) | -26 (48) | 87 (25) | 86 (31) |
| EXOmod | 14 (17) | 17 (15) | 68 (39)* | 67 (46)* | -21 (39)* | -29 (39) | 86 (22) | 84 (25) |
| EXOhigh | 13 (17) | 19 (17) | 69 (42)* | 67 (46)* | -21 (35)* | -20 (49) | 88 (25) | 85 (25) |

#### Subjective feedback

Physical demand (χ2=14.72, *p*<.001), perceived change (χ2=24.22, *p*<.001) and comfort (χ2=13.42, *p*=.001) differed significantly depending on the exoskeleton support level. Post-hoc tests revealed a higher physical demand in EXOdis compared with both EXOmod (Z=3.087, *p*=.006) and EXOhigh (Z=3.344, *p*=.003), whereas EXOmod and EXOhigh (Z=0.257, *p*>.999) did not differ significantly. Perceived change also significantly increased in EXOhigh compared with EXOdis (Z=4.802, *p*<.001) and EXOmod (Z=2.658, *p*=.024). However, there was no statistically significant difference in perceived change between EXOdis and EXOmod (Z=2.144, *p*=.096).

Comfort scores did not differ between EXOdis and EXOmod (Z=1.200, *p*=.690) or EXOdis and EXOhigh (Z=1.972, *p*=.146). However, comfort scores were significantly lower in EXOhigh compared with EXOmod (Z=3.173, *p*=.005). The complete descriptive statistics (n= 17) of the physical demand, perceived change and comfort questionnaire are presented in Figure 5.

**Figure 5.**
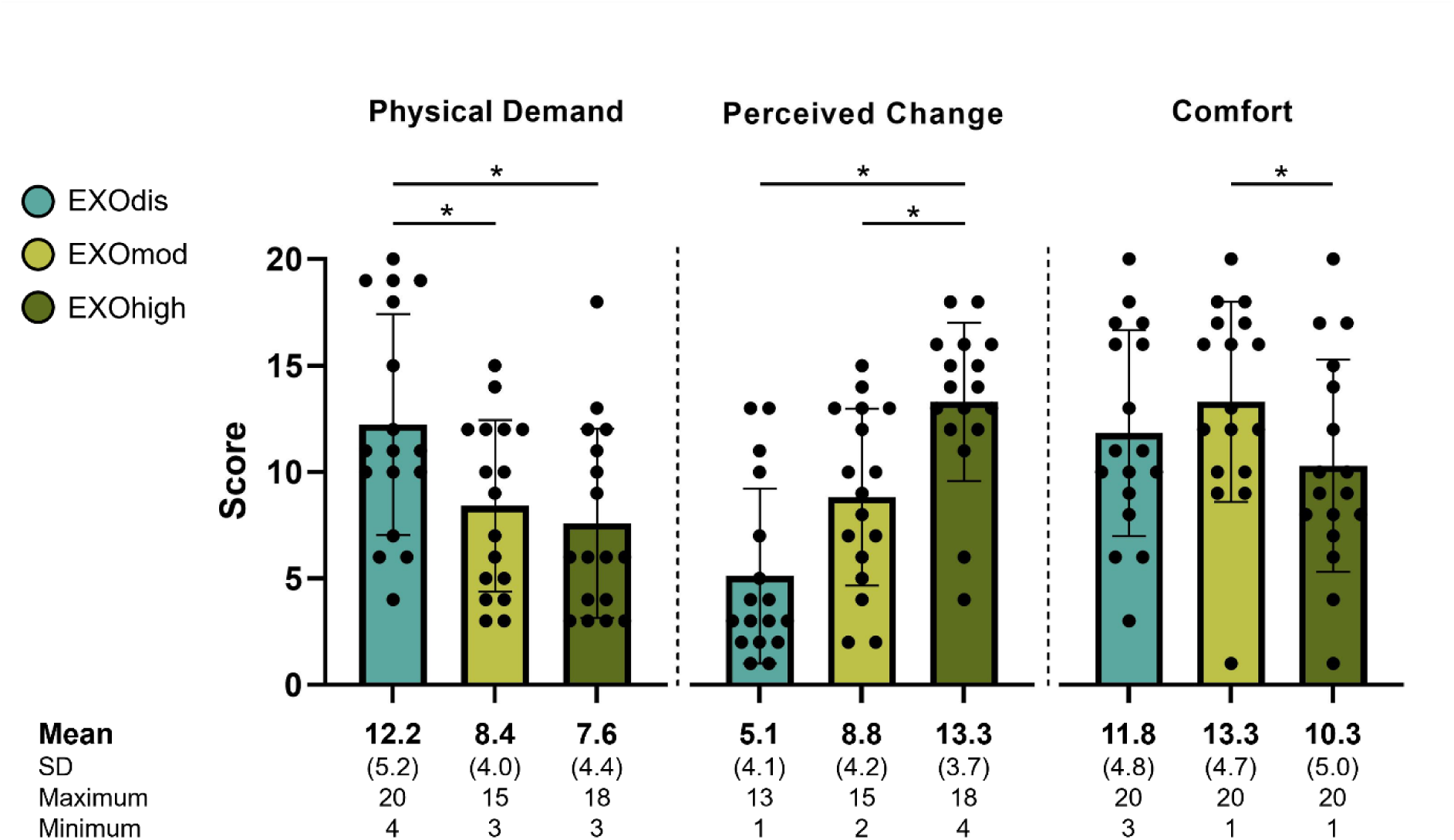
Subjective feedback (n = 17) on the different exoskeleton support levels. The physical demand was perceived to be higher with the exoskeleton support disengaged. Additionally, wearing the exoskeleton was only moderately comfortable, and comfort was reduced with high (EXOhigh) compared with moderate (EXOmod) support. *indicates significant differences between the conditions indicated by the horizontal bars.

## Discussion

The present study investigated whether and how an elastomer-based upper-limb exoskeleton affects posture, movement and muscle activity during two working tasks covering a large shoulder workspace. Furthermore, we evaluated the effect of different levels of exoskeleton support and obtained user feedback on exoskeleton use. We found that exoskeleton use reduced shoulder muscle activity, but only for upper limb postures with an abducted shoulder or extended elbow. Exoskeleton use also modestly altered shoulder postures, with increasing support levels increasing the postural deviation from no exoskeleton use. During the dynamic work task, high exoskeleton support reduced deltoid muscle activity without inducing adverse effects on back muscles. Lastly, subjects reported that the highest exoskeleton support level reduced physical demand and increased perceived change, but perceived comfort was only moderate.

The analysis of muscle activity among different postures during the static task (STA) provided a novel and detailed evaluation of exoskeleton support. A significant muscle activity and posture interaction effect (Fig. 2 & 3) indicates that exoskeleton support was posture specific, with a −2s% to −30s% reductions in muscle activity for static postures with an abducted shoulder or extended elbow (Positions X-I, X-III, and Z-III in Fig. 2). The muscle activity reductions we observed at abducted shoulder postures generally agree with other studies. For example, Maurice et al. (2019) and Van Engelhoven et al. (2018) reported reduced anterior deltoid activity of 24% to 80% during overhead work. Although we found smaller muscle activity reductions, this is likely to be due to the maximum support of EXOhigh. As the EXOhigh support did not match 100% of the calculated arm weight in half of the participants (see *Experimental Protocol*), exoskeleton effects in overhead postures in our study were likely reduced.

Exoskeleton effects were not just limited to overhead postures over the large shoulder workspace we tested in our study. We also found benefits of exoskeleton support at postures with the arm below the horizontal, particularly in extended elbow positions with the arm away from the core (positions Y-III and Z-III). However, muscle activities remained similar between the reference and exoskeleton conditions among positions closer to the core (Fig. 2&3). This finding aligns with De Vries et al. (2019) and Kim et al. (2018), who reported different shoulder muscle activity reductions depending on the work height, with reduced exoskeleton benefits for arm elevation angles below 30 degrees. Accordingly, exoskeleton use has benefits beyond overhead postures, but the benefits are progressively reduced as the arm moves closer to the core.

Muscle activity reductions during exoskeleton use were accompanied by altered upper limb joint angles. However, joint angle differences were rather small (−12° to 5°) and the pathlength analysis revealed that participants’ movements were unaffected by exoskeleton use. This finding supports results by Maurice et al. (2019), who reported no changes of trajectory and velocity during power drill use with exoskeleton support. Similarly, Spada et al. (2018) reported an increased precision during task execution with exoskeleton use. These findings suggest that exoskeleton use does not affect movement patterns during simple work tasks.

During DYN, exoskeleton support significantly decreased AD and MD muscle activities. This is in support of our hypothesis that exoskeleton support is maintained during dynamic movements that cover a large shoulder workspace. In contrast, overall TRAP activity remained similar as it was reduced in overhead postures only (Fig. 3). Taken together, the overall reduction in muscle activity we observed was smaller compared with studies that assessed smaller overhead shoulder workspaces (Schmalz et al., 2019; Van Engelhoven et al., 2018). In addition to the larger workspace tested in the current study, the DYN task itself might have contributed to smaller muscle activity reductions. This is because the DYN task involved raising and lowering of the arm, thereby naturally providing phases with higher and lower shoulder muscle activity, respectively. This alternating on-off muscle activity pattern likely induced less fatigue compared with static overhead tasks requiring constant muscle activity. However, as our study was designed to mimic working tasks over a large shoulder workspace, our findings provide more realistic estimates regarding the benefits of exoskeleton use for work tasks with varying overhead and non-overhead postures.

The absent changes in back muscle activity we observed during exoskeleton use strengthen the case for exoskeleton use during work tasks with a large shoulder workspace. Contrary to our expectations, these findings suggest that the exoskeleton did not introduce unwanted resistance during dynamic movements and that the redirection of forces caused no adverse effects. This result adds new insight to previous findings (De Bock et al., 2021; Huysamen et al., 2018; Maurice et al., 2019) that showed no adverse effects of exoskeleton use on back muscles during overhead tasks, expanding these findings to dynamic tasks over a large shoulder workspace.

Apart from physiological measures, exoskeleton use affected the participants’ subjective feedback. Perception of change was largest for EXOhigh and lowest during EXOdis. Our results further indicate that just wearing the system already led to a moderate perceived change (Fig. 5). However, perceived change merely reflects the perception, without providing insight into whether the change is positive or negative. Therefore, it remains unclear whether the participants perceived the exoskeleton positively or negatively, and whether exoskeleton use induced unintended changes or deliberate (i.e. participant driven) changes in joint kinematics. However, interviews by Maurice et al. (2019) suggest that changes in joint kinematics were, in fact, deliberate. Additionally, our results showed that the support level affected the user’s comfort. The highest exoskeleton support level was perceived as the least comfortable (Fig. 5), which agrees with findings by Van Engelhoven et al. (2018), but somewhat disagrees with the other subjective feedback; the highest support level also reduced the participants’ subjective physical demand and increased their perceived change. Therefore, user comfort should be considered before exoskeleton implementation in real-world workplaces (Gull et al., 2020; Maurice et al., 2019; Smets, 2019), especially because acceptance not only depends on biomechanical benefits, but also on user experience (Bornmann et al., 2020; Siedl & Mara, 2021). Overall, our findings based on subjective feedback highlight that the user-exoskeleton interaction requires optimization.

### Limitations

Although our study demonstrated that positive effects of the tested upper-limb exoskeleton persist over a large shoulder workspace, several limitations need to be considered when interpreting our findings. A key limitation was the inability to match the exoskeleton support to the arm weight of nine participants. Although the reduced support might have attenuated potentially positive exoskeleton effects, these settings reflect the real-world constraints of the tested exoskeleton. Another limitation is related to our EMG analysis of a relatively short time window, which might not accurately represent the muscle activity over the entire task. Further, our evaluation did not include kinetic measurements and included EMG and kinematic measurements only. Lastly, the participants tested do not represent individuals from a typical overhead workforce as they were relatively young and had no professional work experience.

### Conclusion, implication and outlook

This research evaluated the effects of an industrial upper-limb exoskeleton during static and dynamic work tasks over a large shoulder workspace, which prior to this work, were largely unknown. We found that exoskeleton use reduced overall muscle activity beyond overhead postures, but the support was limited to postures with an abducted shoulder or extended elbow. Accordingly, exoskeleton support depends on specific postures and and cannot be generalized over the entire workspace (Kim & Nussbaum, 2019; Musso et al., 2024). Further, the beneficial effects of exoskeleton use depended on the exoskeleton’s support level, which was limited in terms of full arm weight support. Lastly, the exoskeleton’s support level interacted with the subjective perception of change and the perceived comfort, which should be investigated further in future work to help optimize the user-exoskeleton interaction.

Apart from the physiological and subjective effects we found, our evaluation approach of combining static and dynamic standardized work tasks and muscle activity heatmaps might serve as a future standard procedure for upper-limb exoskeleton evaluation (Nussbaum et al., 2019). This approach is simple to implement and can help manufacturers to optimize their systems in terms of posture-specific support and inform customers about potential benefits that can be expected from exoskeleton use. However, future work could expand on our frontal plane heatmaps and incorporate sagittal plane exoskeleton support (De Vries et al., 2019) into a three-dimensional support profile, which might additionally consider kinetic measurements. Finally, future work should evaluate potential long-term effects of exoskeleton use in terms of the prevention of MSDs and potential physiological adaptations to the neuromusculoskeletal system.

## Additional information

### Author contributions

All authors contributed to conception and design of the study and interpretation of data; L.L acquired the data and L.L, B.J.R and P.T contributed to the analysis of data; L.L drafted the manuscript; all authors contributed to revising the manuscript critically for important intellectual content and approved the final version of the manuscript. All authors agree to be held accountable for all aspects of the work, those designated as authors qualify for authorship, and those who qualify for authorship are listed.

## Acknowledgements

The authors thank *Ottobock SE & Co. KGaA* for lending us the exoskeleton used in this study. The company had no influence on the experimental design, data analysis or interpretation and reporting of results.

## References

Barthelme, J., Sauter, M., Mueller, C., & Liebers, F. (2021). Association between working in awkward postures, in particular overhead work, and pain in the shoulder region in the context of the 2018 BIBB/BAuA Employment Survey. BMC Musculoskeletal Disorders, 22(1), 1–12.

Bornmann, J., Schirrmeister, B., Parth, T., & Gonzalez-Vargas, J. (2020). Comprehensive development, implementation and evaluation of industrial exoskeletons. Current Directions in Biomedical Engineering, 6(2).

De Bock, S., Ghillebert, J., Govaerts, R., Elprama, S. A., Marusic, U., Serrien, B., … & De Pauw, K. (2021). Passive shoulder exoskeletons: more effective in the lab than in the field?. IEEE Transactions on Neural Systems and Rehabilitation Engineering, 29, 173–183.

De Leva, P. (1996). Adjustments to Zatsiorsky-Seluyanov’s segment inertia parameters. Journal of biomechanics, 29(9), 1223–1230.

De Vries, A., & De Looze, M. (2019). The effect of arm support exoskeletons in realistic work activities: a review study. J. Ergon, 9(4), 1–9.

De Vries, A., Murphy, M., Könemann, R., Kingma, I., & de Looze, M. (2019). The amount of support provided by a passive arm support exoskeleton in a range of elevated arm postures. IISE Transactions on Occupational Ergonomics and Human Factors, 7(3-4), 311–321.

Dunn, O. J. (1964). Multiple comparisons using rank sums. Technometrics, 6(3), 241–252.

German Federal Ministry of Labor and Social Affairs / Bundesministerium für Arbeit und Soziales (2019). Sicherheit und Gesundheit bei der Arbeit - Berichtsjahr 2018 - Unfallverhütungsbericht Arbeit [Safety and Health at Work - Reporting Year 2018 - Report on Accident Prevention at Work]. Dortmund/Berlin/Dresden: Bundesministerium für Arbeit und Soziales (BMAS), Bundesanstalt für Arbeitsschutz und Arbeitsmedizin (BAuA). Available online: https://www.baua.de/DE/Angebote/Publikationen/Berichte/Suga-2018.html (last accessed on 26.02.2025)

Gull, M. A., Bai, S., & Bak, T. (2020). A review on design of upper limb exoskeletons. Robotics, 9(1), 16.

Hefferle, M., Snell, M., & Kluth, K. (2021). Influence of two industrial overhead exoskeletons on perceived strain–A field study in the automotive industry. In Advances in Human Factors in Robots, Drones and Unmanned Systems: Proceedings of the AHFE 2020 Virtual Conference on Human Factors in Robots, Drones and Unmanned Systems, July 16-20, 2020, USA (pp. 94–100). Springer International Publishing.

Hermens, H. J., Freriks, B., Merletti, R., Stegeman, D., Blok, J., Rau, G., … & Hägg, G. (1999). European recommendations for surface electromyography. Roessingh research and development, 8(2), 13–54.

Holm, S. (1979). A simple sequentially rejective multiple test procedure. Scandinavian journal of statistics, 65–70.

Huysamen, K., Bosch, T., de Looze, M., Stadler, K. S., Graf, E., & O’Sullivan, L. W. (2018). Evaluation of a passive exoskeleton for static upper limb activities. Applied ergonomics, 70, 148–155.

Kim, S., Nussbaum, M. A., Mokhlespour Esfahani, M. I., Alemi, M. M., Alabdulkarim, S., & Rashedi, E. (2018). Assessing the influence of a passive, upper extremity exoskeletal vest for tasks requiring arm elevation: Part I - “Expected” effects on discomfort, shoulder muscle activity, and work task performance. Applied Ergonomics, 70, 315–322.

Kim, S., & Nussbaum, M. A. (2019). A follow-up study of the effects of an arm support exoskeleton on physical demands and task performance during simulated overhead work. IISE Transactions on Occupational Ergonomics and Human Factors, 7(3-4), 163–174.

Kim, S., Nussbaum, M. A., Smets, M., & Ranganathan, S. (2021). Effects of an arm-support exoskeleton on perceived work intensity and musculoskeletal discomfort: An 18-month field study in automotive assembly. American Journal of Industrial Medicine, 64(11), 905–914.

Knight, J. F., Baber, C., Schwirtz, A., & Bristow, H. W. (2002, October). The Comfort Assessment of Wearable Computers. In iswc (Vol. 2, pp. 65–74).

Maurice, P., Čamernik, J., Gorjan, D., Schirrmeister, B., Bornmann, J., Tagliapietra, L., … & Babič, J. (2019). Objective and subjective effects of a passive exoskeleton on overhead work. IEEE Transactions on Neural Systems and Rehabilitation Engineering, 28(1), 152–164.

Musso, M., Oliveira, A. S., & Bai, S. (2024). Influence of an upper limb exoskeleton on muscle activity during various construction and manufacturing tasks. Applied Ergonomics, 114, 104158.

Nussbaum, M. A., Lowe, B. D., de Looze, M., Harris-Adamson, C., & Smets, M. (2019). An introduction to the special issue on occupational exoskeletons. IISE Transactions on Occupational Ergonomics and Human Factors, 7(3-4), 153–162.

Nuzzo, R. (2018). Percent differences: Another look. PM&R, 10(6), 661–664.

Otten, B. M., Weidner, R., & Argubi-Wollesen, A. (2018). Evaluation of a novel active exoskeleton for tasks at or above head level. IEEE Robotics and Automation Letters, 3(3), 2408–2415.

Ottobock SE & Co. KGaA. (2019). *16ES100=2 Paexo Shoulder: Original instructions*. 37115 Duderstadt, Germany: Ottobock.

Schmalz, T., Schändlinger, J., Schuler, M., Bornmann, J., Schirrmeister, B., Kannenberg, A., & Ernst, M. (2019). Biomechanical and Metabolic Effectiveness of an Industrial Exoskeleton for Overhead Work. International Journal of Environmental Research and Public Health, 16(23). 4792.

Siedl, S. M., & Mara, M. (2021). Exoskeleton acceptance and its relationship to self-efficacy enhancement, perceived usefulness, and physical relief: A field study among logistics workers. Wearable Technologies, 2, e10.

Smets, M. (2019). A field evaluation of arm-support exoskeletons for overhead work applications in automotive assembly. IISE Transactions on Occupational Ergonomics and Human Factors, 7(3-4), 192–198.

Spada, S., Ghibaudo, L., Gilotta, S., Gastaldi, L., & Cavatorta, M. P. (2018). Analysis of exoskeleton introduction in industrial reality: main issues and EAWS risk assessment. In Advances in Physical Ergonomics and Human Factors: Proceedings of the AHFE 2017 International Conference on Physical Ergonomics and Human Factors, July 17-21, 2017, The Westin Bonaventure Hotel, Los Angeles, California, USA 8 (pp. 236–244). Springer International Publishing.

Van Engelhoven, L., & Kazerooni, H. (2019, March). Design and intended use of a passive actuation strategy for a shoulder supporting exoskeleton. In 2019 Wearable Robotics Association Conference (WearRAcon) (pp. 7–12). IEEE.

Van Engelhoven, L., Poon, N., Kazerooni, H., Barr, A., Rempel, D., & Harris-Adamson, C. (2018, September). Evaluation of an adjustable support shoulder exoskeleton on static and dynamic overhead tasks. In Proceedings of the Human Factors and Ergonomics Society Annual Meeting (Vol. 62, No. 1, pp. 804–808). Sage CA: Los Angeles, CA: SAGE Publications.

Vicon Motion Systems. (2023). Plug-in Gait reference guide. Vicon Motion Systems Ltd. Available online: https://help.vicon.com/download/attachments/11378719/Plug-in%20Gait%20Reference%20Guide.pdf (last accessed on 26.02.2025)

