## Supplementary material for "A passive upper-limb exoskeleton effectively reduces shoulder muscle activity over a large shoulder workspace"

#### Title page

#### Title


#### Author names & affiliations

Leon Lauret[1], Brent James Raiteri[1,2], Paolo Tecchio[1], Daniel Hahn[1,2]

[1] Human Movement Science, Faculty of Sport Science, Ruhr University Bochum, Bochum, North Rhine-Westphalia, Germany

[2] School of Human Movement and Nutrition Sciences, The University of Queensland, Brisbane, Queensland, Australia

#### Corresponding author

Leon Lauret

Gesundheitscampus Nord 10, Bochum, North Rhine-Westphalia, 44801, Germany

### Supplementary Material

#### Data availability

The collected raw data and scripts used in this study will be made available upon publication. Links to the raw data and scripts will be shared upon publication.

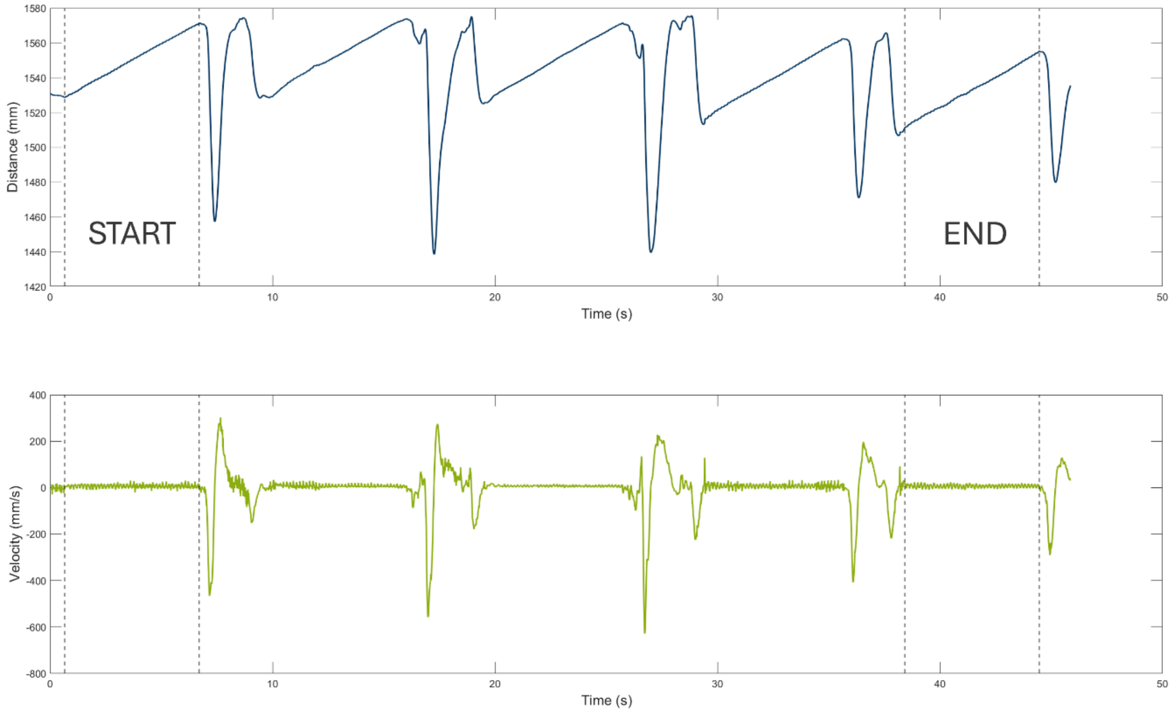


**Supplementary Figure 1. Methodology used to identify START (i.e. the first screw) and END (i.e. the last screw) during DYN.** During DYN, kinematics and muscle activities were recorded for the first five and last five screws for each support level. The figure shows representative data of a five-screw recording during DYN with displacement (blue line) and velocity (green line) of the right finger marker in the driving direction over time (s). To identify the first (START) and identical last screw (END) from the two five-screw recordings, manual and subjective identification was used. The beginning of START was defined as the onset of the first positive linear displacement with constant velocity (first vertical dashed line), whereas the end of START was defined as the first deviation from constant velocity (second vertical dashed line). END was defined from recordings of the last five screws in a similar manner, with the on- and offset of END defined identically as for START but for the fifth screw. Pathlength was calculated over the first five and last five screws.

#### QUESTIONNAIRE FOR SUBJECTIVE FEEDBACK ON THE EXOSKELETON

Subject / Condition:______________________

***Please mark each scale at the point that best indicates your experience with the exoskeleton with the help of the questions asked in the description box from low to high***

Physical Demand

Low — High

How much physical activity was required?
Was the task easy or demanding?

Perceived change

Low — High

Does the exoskeleton make you feel physically different?

Comfort

Low — High

Does using the exoskeleton make you feel (un-)comfortable?

**Supplementary Figure 2. The questionnaire used to assess the subjective feedback on wearing the exoskeleton** during DYN in terms of physical demand of the task, perceived change and comfort of the exoskeleton. The questionnaire used a 21 point scale, with scores from 0 (“low”) on the left to 20 (“high”) on the right.
